# Quantifying PD1 saturation by PDL1 in tumor tissue using a novel RNA aptamer-based assay

**DOI:** 10.64898/2026.04.06.716702

**Authors:** Suresh Veeramani, Chaobo Yin, Nanmeng Yu, Kristen Coleman, Brian Smith, George J. Weiner

## Abstract

**Background:** Therapeutic agents targeting the PD1-PDL1 interaction are of great clinical value, however accurately predicting which patients are most likely to benefit is challenging. Improved predictive biomarkers for anti-PD1 therapy are clearly needed. Quantifying PD1 saturation by PDL1 in tumor tissue has the potential to serve as such a biomarker. Here we report a novel bioassay called the PD1 Ligand Receptor Complex Aptamer (LIRECAP) assay and demonstrate it can be used to quantify the saturation of PD1 by PDL1 in formalin-fixed paraffin-embedded tumor biospecimens.

**Results:** The PD1 LIRECAP assay was developed by identifying a pair of RNA aptamers. One aptamer preferentially binds to unoccupied PD1 (P aptamer) and the other to the PD1-PDL1 complex (C aptamer). P and C aptamers were added together to a formalin-fixed sample, and bound aptamer extracted. A 2-color qRT-PCR assay using a single set of primers was used to determine the ratio of the sample-bound C to P aptamers (C:P ratio) which reflected PD1 saturation by PDL1 in the sample. Quantification of PD1 saturation by PDL1 as determined by the PD1 LIRECAP assay correlated closely with PD1-mediated signaling and PD1-PDL1 proximity. Analysis of sarcoma FFPE biospecimens confirmed the assay is technically reproducible on clinical biospecimens. There were significant differences in PD1 saturation by PDL1 between patients as well as considerable intratumoral heterogeneity.

**Conclusions:** The PD1 LIRECAP assay is novel assay that can be used to quantify PD1 saturation by PDL1 in clinical biospecimens. The assay is technically feasible, reproducible, and has the potential to serve as a superior predictive biomarker for PD1/PDL1-based therapy. Similar assays based on this platform could be used in other systems and settings to quantify interaction between two molecules.

## Background

Ligand-receptor interactions play a central role in biology and therapeutics; however, quantification of ligand-receptor interactions is understudied for its biomarker potential. Established methods, such as colocalization, fluorescence resonance energy transfer (FRET) and proximity ligation assays (PLAs) rely on the evaluation of the receptor-ligand proximity[1–3], not on direct measurement of interaction. These assays are valuable in some research settings but are semi-quantitative and can be technically challenging to perform, particularly at high-throughput volumes[4–6].

Our group previously developed a novel RNA aptamer-based approach to quantify the saturation of a circulating receptor by its ligand[7, 8] termed the Ligand-Receptor Complex-binding Aptamer (LIRECAP) assay. The LIRECAP assay is based on a pair of RNA aptamers with one aptamer preferentially binding to unoccupied receptor and the other preferentially binding to the ligand-receptor complex. These aptamers have distinct central regions but are of the same length and have the same 5’ and 3’ ends. When they are added in equal amounts to a biospecimen, the ratio of bound aptamers reflects the receptor saturation by ligand. The aptamer ratio can be determined through a single, TaqMan-based RT-qPCR assay performed using a single set of primers. This adds a robust level of internal control and provides a quantifiable measure of the saturation of the receptor by its ligand that is independent of the absolute amounts of receptor or ligand in the sample. The LIRECAP assay represents a novel, sensitive and technically straight-forward approach for quantifying the saturation of a receptor by its ligand[8]. To date, it has only been applied in proof-of-concept studies with soluble CD25-IL2 receptor-ligand complexes.

Therapy directed at blocking the interaction between Programmed cell death protein 1 (PD1) and its ligand PDL1 has generated considerable excitement in the field of cancer therapeutics[9–11]. Signaling through PD1 mediated by PDL1 leads to inhibition of TCR signaling and suppresses the anti-tumor activity of tumor specific T cells[12]. Blocking PD1-PDL1 interaction with anti-PD1 or anti-PDL1 monoclonal antibodies, collectively referred to here as PD1 blockade therapy, reverses T cell inhibition and can lead to tumor regression. Unfortunately, all patients treated with PD1 blockade are exposed to potential toxicities while only a minority of patients benefit from such therapy[13, 14]. The ability of current biomarkers, such as T cell infiltration, tumoral PDL1 expression, tumor mutational burden and microsatellite instability, to predict which patients are likely to benefit, is limited[15, 16]. Intratumoral heterogeneity in PD1 and PDL1 levels further complicates biomarker-based clinical decision making[17, 18]. Preliminary studies indicate that studying PD1-PDL1 proximity in biospecimens may predict outcome more effectively than the standard biomarkers[19–22] however, as stated above, they are not well suited to broad clinical application.

Better biomarker assays, including those measure PD1-PDL1 interactions, are clearly needed. Here, we report on the development of a PD1 LIRECAP assay capable of quantifying the saturation of PD1 by PDL1 in formalin-fixed paraffin embedded (FFPE) tumor tissue and discuss its potential as a biomarker for PD1-based therapy.

## Methods

### Reagents

The DNA template for SEL2 RNA library, containing the random N20 central region (SEL2-N20), PCR primers and fluorochrome-labeled TaqMan probes were chemically synthesized (Integrated DNA technologies). 2’ fluoro-modified dUTP and dCTP were purchased from TriLink Biotech. Purified E. coli-expressed recombinant T7 RNA polymerase (Y639F) was used for in vitro transcription of the 2’ fluoro-modified RNA aptamer library[7]. Recombinant human Fc- or Histidine-tagged proteins, including PD1, PDL1, Glyoxalase and Ubiquitin (UbQ), were purchased from Acrobiosystems, Fisher Scientific and R&D Systems. Dynabeads® for binding histidine-tagged or Fc-tagged proteins and RNA/DNA-modifying enzymes, including Taq DNA polymerase, RNase-free DNase I, RNase Inhibitor and RNase A, were purchased from Thermo-Scientific. Protease inhibitors, inorganic pyrophosphatase and nucleotides (NTPs and dNTPs) were purchased from Promega and New England Biolabs. The Naveni PD1/PD-L1 Atto647N Proximity assay ligation kit (Navinci) for detecting PD1-PDL1 proximity was purchased from Sapphire North America. All other chemical and biological reagents were obtained from Sigma Aldrich and Fisher Scientific.

### Cell lines

The Jurkat-Raji PD-1/PD-L1 assay (Bio-IC™) co-culture system used for studying PD1-PDL1 interaction was purchased from Invivogen. The system includes,

1. Raji-APC-hPDL1 (Raji-PDL1) – Raji cells modified to express human PDL1 and a modified MHC,
2. Raji-APC-Null (Raji-Null) – Raji cells that do not express PDL1 and modified MHC, and
3. Jurkat-Lucia™ TCR-hPD-1 (Jurkat-PD1) – Jurkat T cells modified to express human PD1. These cells also express TCR reactive to Raji-APC cells and a luciferase reporter gene responsive to TCR-induced NF-AT signaling.

All cells were cultured in IMDM (Thermo-Fisher) containing 10% FBS (Thermo-Fisher), 2 mM L-Glutamine (Thermo-Fisher), 100 IU/mL Penicillin, 100 ug/mL Streptomycin and 100 ug/mL Normocin (Invivogen). Selection antibiotics, as per the manufacturer’s instructions, were included during every other subculture to maintain the purity of recombinant cells.

### Human sarcoma biospecimens

De-identified FFPE blocks from six patients diagnosed with human sarcoma who provided consent for use of tissue in research through a protocol approved by the University of Iowa Human Subjects Office were obtained from the Biospecimens and Molecular Epidemiology Resource, University of Iowa Holden Comprehensive Cancer Center. All the protocols are approved by the Institutional Review Board of the University of Iowa. Sections of 5 μm-thick were made as slides for IHC and PLA and sections of 10 μm-thick were made as scrolls for the PD1 LIRECAP assay. Select slides were stained for CD3 to confirm T cell infiltration and PDL1 expression. Based on PDL1 expression on tumor cells, sarcomas were classified as PDL1^high^ or PDL1^low^ (Supplementary Table S4).

### Selection of PD1- and PD1-PDL1 Complex-binding aptamer library

Enriched RNA aptamer libraries that preferentially recognized unoccupied human PD1 or PD1-PDL1 complex proteins were selected using a modified SELEX strategy. Briefly, the starting RNA library was transcribed from the SEL2-N20 DNA template to incorporate 2’F-modified pyrimidines. One nmole of the starting RNA library in LIRECAP buffer (HEPES-buffered saline, pH 7.5, containing 2 mM CaCl_2_ and MgCl_2_) was folded at 75^0^C for 10 minutes followed by cooling to 37^0^C. The folded library was diluted to 2 μm concentration. SELEX conditions used for pre-clearing non-specific aptamers and selecting target-specific aptamers are summarized in Supplementary Table S1. All the SELEX incubations were done on 4^0^C to minimize dissociation of PD1-PDL1 complexes. Bound aptamers were extracted using phenol-chloroform and amplified by RT-PCR using the SEL2 primers. Amplified DNA was transcribed into 2’ fluoro-modified RNA for the next SELEX round. After round 5, aptamer pools underwent a post-clearing step to minimize PDL1-binding aptamers from both libraries and PD1-binding aptamers from the Complex library. A total of eight rounds of SELEX per target were performed.

#### Bioinformatics analysis

RNA aptamer pools and corresponding PCR DNA from each SELEX round was sequenced by Illumina-based next-generation sequencing (Genomics Division, University of Iowa), as described[7]. Individual aptamer sequences from each round were analyzed using a standardized aptamer bioinformatics workflow[7, 23, 24]. Briefly, a persistence filter and an abundance filter were first applied to the large dataset with all the reads. For this, RNA aptamers that were found in at least three of eight SELEX rounds (Persistence filter) as well as that had at least 30 read counts in any of the eight SELEX rounds (Abundance filter) were chosen. Based on these initial criteria, a total of 5,401 P aptamers and 4,625 C aptamers were identified. The final list of potential candidate sequences was selected based on three specific criteria:

1. <u>Enrichment in the final library</u>: Candidates with high fold enrichment in round 8 compared to the starting library (round 0), indicating high target binding.
2. <u>Progressive enrichment</u>: Sequences showing increasing enrichment across successive rounds of SELEX, demonstrating a gradual increase in binding.
3. <u>Target-specific enrichment</u>: Finally, aptamers were identified that showed higher enrichment in the relevant SELEX library (e.g., higher enrichment for P aptamers in PD1 SELEX than in Complex SELEX and vice versa). This step ensured that sequences that were enriched in both pools, indicative of highly cross-reactive aptamers, were excluded.

### Aptamer specificity assay

Dynabeads were coated with 50 nM recombinant His-tagged human PD1 or His-tagged PDL1. To create beads containing PD1-PDL1 complex, an aliquot of PD1-coated beads was incubated with 200 nM Fc-tagged PDL1. For creating varying PD1 saturation, PD1-coated beads were incubated with a serial dilution of Fc-tagged PDL1. Complex formation was facilitated by incubating for 15 minutes at 37°C followed by 90 minutes at 4°C. After washing off unbound PDL1, PD1- and Complex-coated beads were fixed with 2% formaldehyde for 30 minutes at RT. Formalin-fixed targets were washed extensively with LIRECAP wash buffer (10mM HEPES-buffered saline containing 2 mM CaCl2 and 0.05% Tween-20) to remove traces of formaldehyde. Beads were then blocked with LIRECAP blocking buffer (HEPES-buffered saline, pH 7.5, containing 2 mM CaCl_2_, MgCl_2_, 0.5 mg/ml BSA and tRNA). For evaluation of aptamer binding, target-coated beads were incubated with 100 nM folded aptamer candidates for 90 minutes at 4°C Supplementary Table S2. After washing off unbound aptamers, target-bound aptamers were extracted with phenol-chloroform and quantified using SyBR green-based qPCR. To normalize for differences in RNA extraction, phenol-chloroform spiked with a reference sequence, M12-23, and bound aptamer values were normalized to M12-23 values[7].

#### Cell-based evaluation of PD1 signaling and PD1 saturation by PDL1

The commercially available Jurkat-Lucia™ TCR-hPD-1 cell and Raji cell co-culture system (Invivogen) was used for evaluation of both PD1 signaling and PD1 saturation by PDL1 (Invivogen). Raji cell mixtures containing different PDL1 levels were produced by mixing various ratios of PDL1-negative (Raji-Null) and PDL1-positive (Raji-PDL1) cells (Supplementary Table S3). These Raji cell mixtures were then mixed in a 1:1 ratio with Jurkat-Lucia™ TCR-hPD-1 (Jurkat-PD1) cells. Such an approach of using a consistent Raji to Jurkat ratio (1 to 1) but altering the fraction of Raji cells that expressed PDL1 allowed for adjustment of PD1 saturation by PDL1 (Supplementary Table S3). Cells were co-cultured for 6 hours at 37^0^C. To quantify the PD1 signaling and Jurkat T cell inhibition, conditioned media was harvested and mixed with QUANTI-Luc™ 4 Lucia/Gaussia detection reagent (Invivogen). Bioluminescence was immediately measured using a standard plate reader equipped with filters measuring chemiluminescence. The percent inhibition of Jurkat T cell activation was calculated based on the bioluminescence values of each co-culture normalized to control co-cultures (e.g. no inhibition with Jurkat-PD1 and Raji-Null cells, full inhibition with Jurkat PD1 and Raji-PDL1 cells). After collection of supernatants, cell pellets were formalin fixed at 2% formaldehyde for 30 minutes at room temperature. After extensive washing, samples were blocked overnight using blocking buffer and incubated with 25 nM equimolar mix of P and C aptamers for 90 minutes at 4^0^C. Bound aptamers were extracted using Trizol-based method and quantified by TaqMan-based RT-qPCR to derive the C:P ratio.

#### PD1 LIRECAP assay on FFPE biospecimens

FFPE blocks were prepared from the formalin-fixed Jurkat-Raji cell pellets and the human sarcoma tumors using standard tissue processing protocols[25] by the University of Iowa Holden Comprehensive Cancer Center biorepository and molecular epidemiology resource. Blocks were sectioned into 10 μm-thick scrolls for the PD1 LIRECAP assay. Scrolls were deparaffinized and rehydrated using standard IHC procedures.

The resulting tissue was macerated using a pellet pestle. In some cases, macerated cells were divided into separate aliquots so technical reproducibility could be evaluated by assessing samples from the same tissue on different PD1 LIRECAP assay runs. Samples were then blocked overnight using blocking buffer and incubated with 25 nM equimolar mix of P and C aptamers for 90 minutes at 4^0^C. After extensive washing, bound aptamers were extracted using Trizol-based method. Bound aptamers were quantified using the TaqMan-based RT-qPCR method to derive the C:P ratio.

### Proximity ligation assay

Proximity ligation assays for detecting PD1-PDL1 interaction in the FFPE biospecimens were done using a commercially available Naveni PD1/PD-L1 Atto647N Proximity assay ligation kit (Navinci). Briefly, 5 μm-thick sections were stained per the manufacturer’s recommendation. Slides were imaged using Keyence X-800 confocal microscope and the number of PLA spots (PD1-PDL1 proximity) and DAPI spots (cell nucleus) were counted in multiple areas in each slide. Average PLA positivity was calculated by normalizing the average number of PLA spots to the average number of DAPI-positive cells in each frame and each slide and expressed as PLA positivity per cell or as percent positivity.

### Generation of a standard curve for quantifying PD1 saturation index

The following approach was used for the generation of a standard curve using recombinant human 6X His-tagged PD1 and Fc-tagged PDL1 proteins (Acrobiosystems).

1. *<u>Creation of beads coated with PD1 and various concentrations of PDL1</u>*: PD1-coated beads were created by incubating PD1 with histidine-binding Dynabeads® (Thermo-Fisher). These beads were incubated with varying concentrations of PDL1 for 90 minutes at 4^0^C. PD1 saturation was calculated as percentage using the PD1 and PDL1 concentration as well as their affinity values (Kd = 38.5 nM) (Acrobiosystems). Target-coated beads were formalin-fixed and blocked with cold blocking buffer. PD1 saturation by PDL1 for each sample was calculated using the following quadratic formula that utilizes the Kd value for PD1-PDL1 interaction[26].

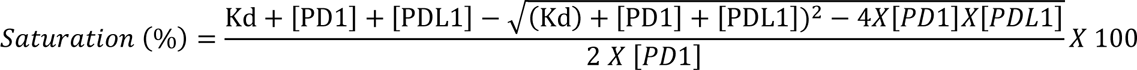
2. *<u>Reference control</u>*: To reduce inter-sample technical variations, equal concentration of Streptavidin-coated MyOne® C1 beads (Thermo-Fisher) coated with Desthiobiotin-labeled M12-23 aptamers was added.
3. *<u>Incubation of aptamers</u>*: Beads containing varying levels of PD1 saturation by PDL1 were incubated with an equimolar mix of P and C aptamers each at 100 nM concentration for 90 minutes at 4^0^C.
4. *<u>Bound aptamer extraction</u>*: Aptamer bound samples are washed extensively to remove unbound aptamers. Bound RNA aptamers and the reference control were extracted using Phenol-chloroform-based method.
5. *<u>Aptamer quantification</u>*: The ratio of bound C to P aptamer was quantified using a TaqMan-based RT-qPCR, a single set of PCR primers and TaqMan probes specific for each aptamer. Reference control in each sample was amplified using a SyBR green-based qPCR using SEL1 primer set[7] and was used to normalize each sample. C:P ratio was then calculated for each sample.
6. *<u>Generation of standard curve</u>*: Standard curve was created by plotting the C:P ratio against the calculated PD1 saturation by PDL1 (%).

### Statistical analysis

All experiments, unless specifically mentioned, were repeated at least twice to ensure reproducibility. Data points from multiple assays, done with similar conditions, were pooled and mean and standard error of mean (SEM) were calculated. Covariance, correlation and significance of the mean (*p* value) were analyzed using the GraphPad Prism v10. A *p*-value of less than 0.05 was considered significant. Linear mixed effects regression was used to estimate intraclass correlation coefficient (ICC) for repeated measurements using R-software (v4.5.1) (Vienna, Austria). For technical reproducibility and intratumoral heterogeneity analyses, random effects were included for experimental replicates. Estimates are reported along with bootstrapped 95% confidence intervals.

## Results

### Selection of aptamers for the PD1 LIRECAP assay

RNA aptamers that preferentially bind to unoccupied PD1 or to the PD1-PDL1 complex were selected using the Systematic Evolution of Ligands by Exponential enrichment (SELEX) approach using recombinant PD1 and PDL1 (Schema in Fig. 1A)[27]. Briefly, the starting RNA aptamer library (Rd 0) was divided into two equal aliquots – one for selecting PD1-binding aptamers (P aptamer pool) and the other for selecting PD1-PDL1 complex-binding aptamers (C aptamer pool). A total of eight selection rounds, consisting of both negative and positive selection, were performed for each pool (Supplementary Table S1). Assessment of target binding of P and C aptamer pools after round 3 and round 7 revealed progressive enrichment for aptamers binding to PD1 or the PD1-PDL1 Complex respectively (Figs. 1B and 1C). Sequencing of aptamer pools indicated a steady decline in the number of unique aptamer sequences in both the P and C aptamer pools after each round of selection (Supplementary Fig. S1). Enrichment was most notable between first and fifth SELEX rounds with limited enrichment occurring in the later rounds (Fig. 1D).

**Fig. 1:**
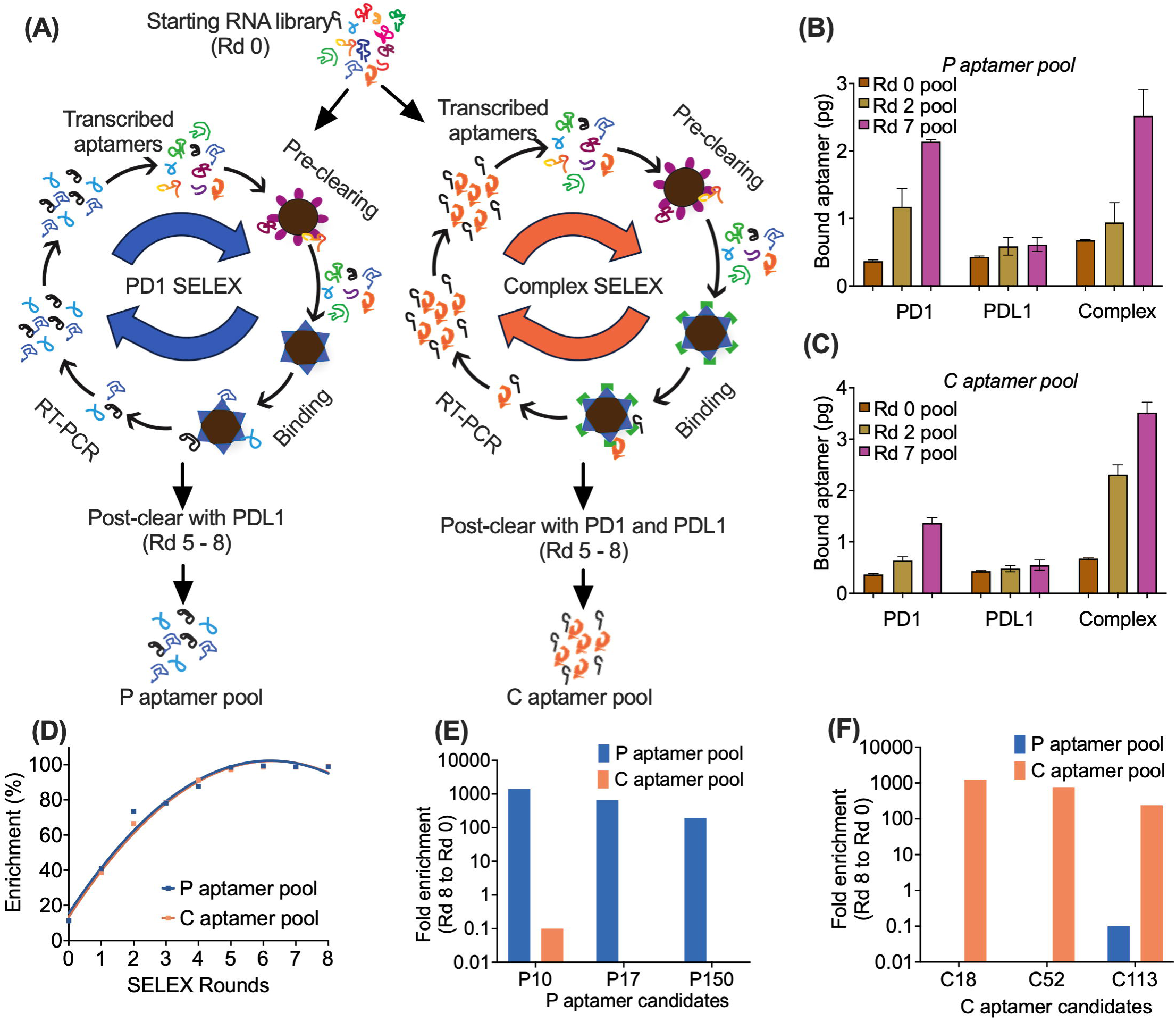
Selection of lead aptamer candidates for PD1 LIRECAP assay. (A) Schematic illustration of the SELEX approach used to generate enriched aptamer pools recognizing unoccupied PD1 (PD1 SELEX) and PD1-PDL1 Complex (Complex SELEX). (B) Binding specificity of P aptamer pools to recombinant PD1, PDL1 and PD1-PDL1 Complex. (C) Binding specificity of enriched C aptamer pools to recombinant PD1, PDL1 and PD1-PDL1 Complex. Data shown is a representative of two independent experiments. (D) Percent enrichment of unique aptamer sequences in P and C aptamer pools following various rounds of selection as determined by Illumina-based high-throughput sequencing. (E) Fold enrichment of lead P aptamer candidates (P10, P17 and P150) based on copy number in the final enriched pool normalized to the starting library (Rd 0). (F) Fold enrichment of lead C aptamer candidates (C18, C52 and C113) based on copy number in the final enriched pool normalized to the starting library (Rd 0).

Sequences were identified as candidate aptamers that (1) showed sequential enrichment as SELEX rounds progressed and (2) were predominantly in either the P aptamer pool or the C aptamer pool, but not both. The final determination of which aptamer sequences to synthesize and evaluate was made using the bioinformatics workflow and filters described in the Methods section. Three candidate aptamers from the P aptamer pool (designated as P10, P17 and P150) and three from C aptamer pool (designated C18, C52 and C113) were chosen for further analysis. The frequency of these aptamers in the final P aptamer pool and C aptamer pool is shown in Figs. 1E, 1F and Supplementary Table S2.

### Evaluation of lead P and C aptamers

Lead candidates were transcribed into 2’-fluoro-modified RNA and evaluated for target specificity using magnetic beads coated with formalin-fixed PD1, PDL1 or the PD1-PDL1 Complex. Formalin fixation was done for three reasons: First, it stabilized the PD1-PDL1 complex. Second, it assured that binding of the aptamers would not interfere with the PD1-PDL1 interaction. Third, it assured aptamers could bind to formalin fixation targets since the intent was to use these aptamers to assess PD1 saturation by PDL1 in clinical FFPE tissue. P aptamers (P10, P17, and P150) bound more to unoccupied PD1 than to PD1-PDL1 complex or to unoccupied PDL1 (Figs. 2A-2C). C aptamers (C18, C52, and C113) bound more to PD1-PDL1 complex than to unoccupied PD1 or unoccupied PDL1 (Figs. 2D-2F). Since PD1 can also engage with PDL2, we evaluated the binding patterns of P17 and C18 aptamers to the PD1-PDL2 complex. The aptamers displayed comparable binding to the PD1-PDL2 complex as they did to the PD1-PDL1 complex, which implies potential cross-reactivity or recognition of shared structural features in PD1-mediated interactions (Supplementary Figs. S2A and S2B).

**Fig. 2:**
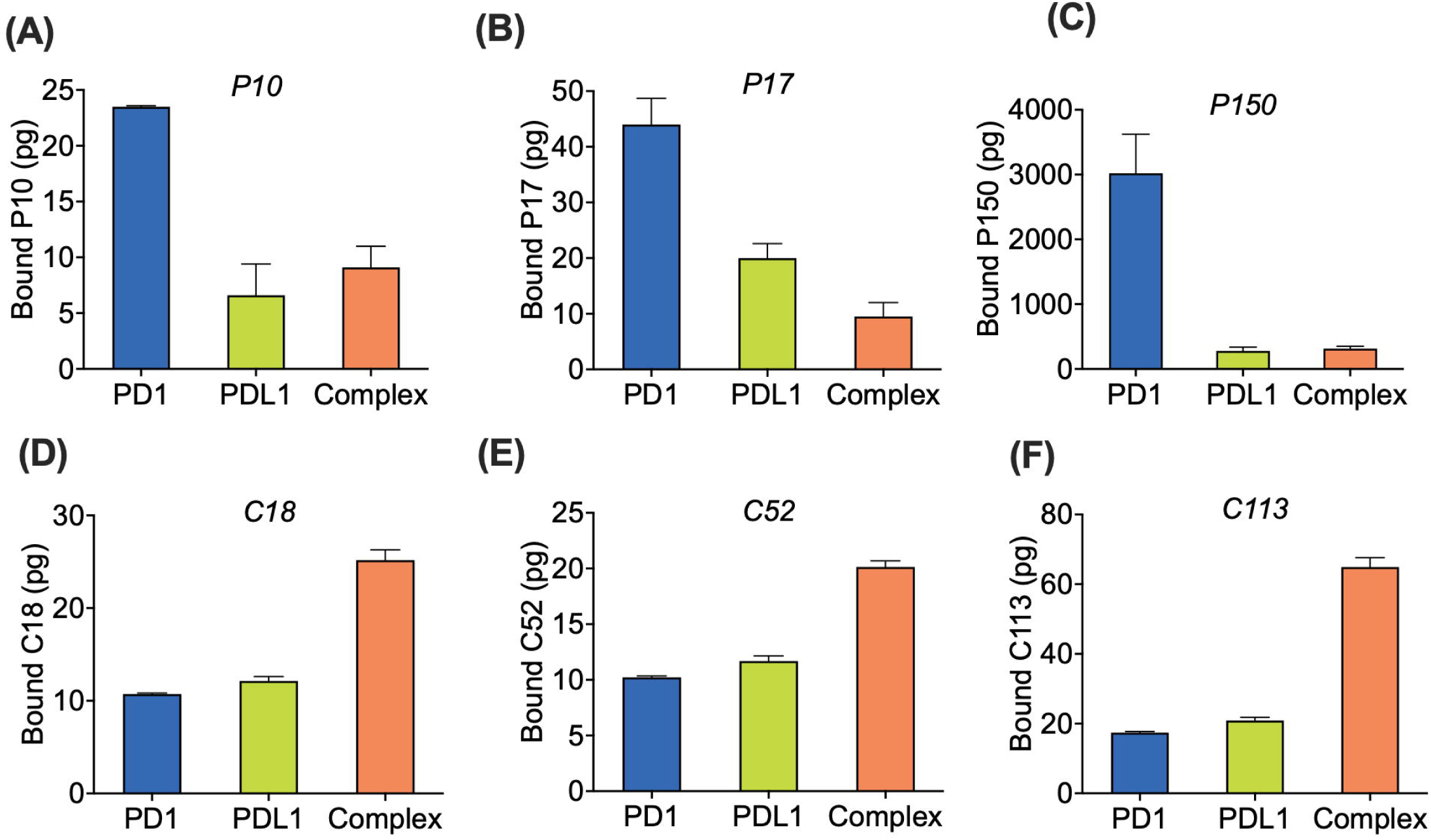
Specificity of lead P and C aptamer candidates towards PD1, PDL1 and PD1-PDL1 complex. Lead P aptamer candidates (A) P10, (B) P17, and (C) P150 binding to unoccupied PD1, PDL1 and PD1-PDL1 Complex. Lead C aptamer candidates (D) C18, (E) C52, and (F) C113 binding to PD1-PDL1 Complex, unoccupied PD1 and PDL1. Data are representative of two independent experiments.

P17 and C18 were chosen for development of the PD1 LIRECAP assay because they bound to formalin-fixed targets and did not cross compete for target binding (Supplementary Fig. S3A). Unique TaqMan probes were synthesized for P17 and C18 that bound to their respective variable regions (Fig. S3B and S3C). TaqMan-based qPCR analysis, using primers shared by both aptamers, demonstrated P17 and C18 have similar amplification efficiency (PCR efficiency P17 = 97.3%; C18 = 98.5%).

### Bound aptamer ratio correlates with PD1 signaling

The ratio of bound C18 to P17 (C:P ratio), as determined by the PD1 LIRECAP assay (schema in Fig. 3A), was evaluated using the Jurkat-Lucia™ TCR-hPD-1 cell and Raji cell co-culture system (Invivogen) that also allows for evaluation of PD1 signaling. This system consists of PDL1-negative (Raji-Null), PDL1-positive (Raji-PDL1) Raji cells and Jurkat-Lucia™ TCR-hPD-1 cells (Jurkat-PD1) containing a modified TCR that recognizes MHC-peptide on Raji cells and leads to TCR activation. TCR activation in the Jurkat-PD1 is visualized using the Lucia luciferase reporter gene.

**Fig. 3:**
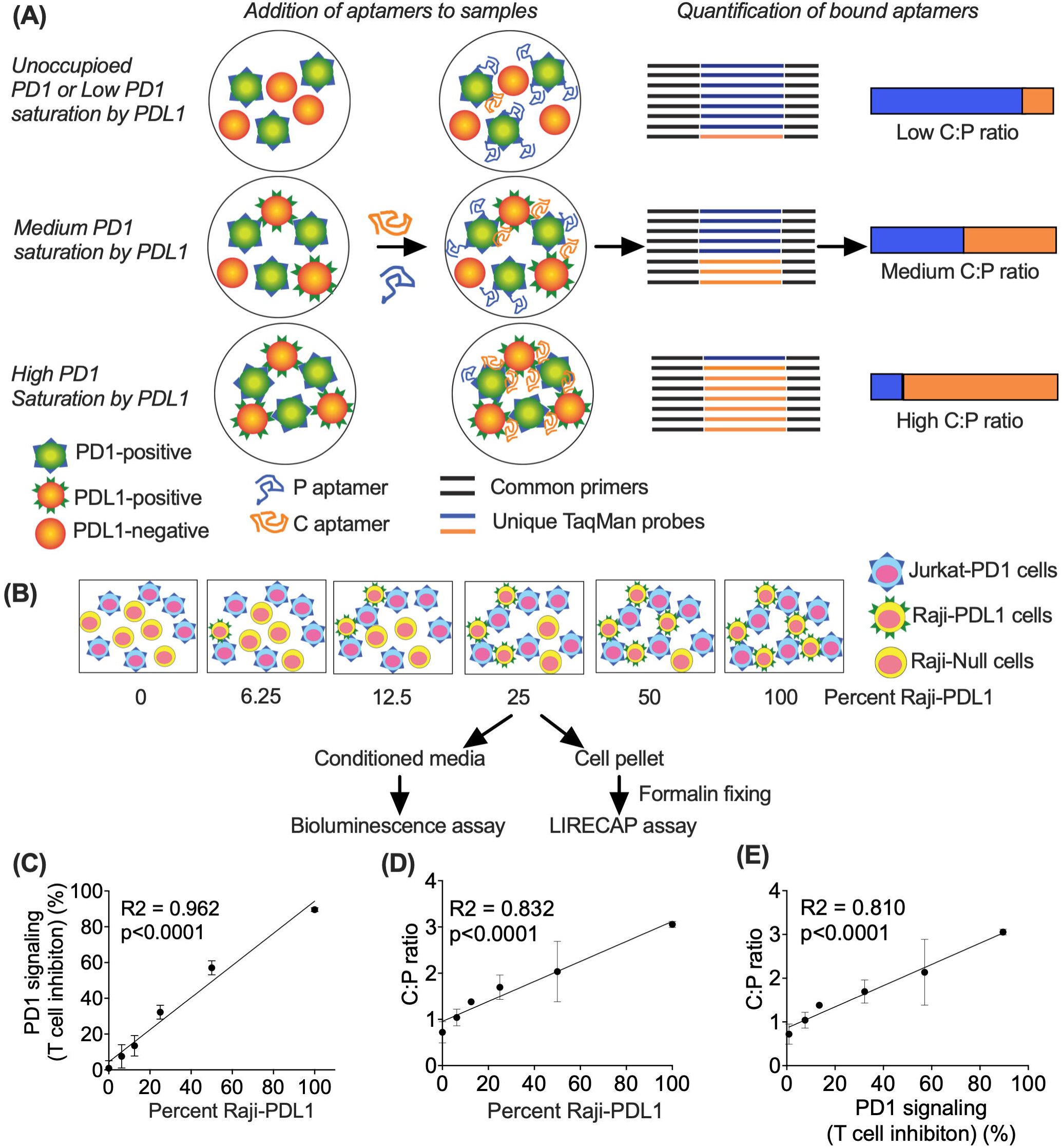
Correlation between PD1 LIRECAP assay (C:P aptamer ratio) and PD1 signaling in the Jurkat-Raji co-culture system. (A) Schematic illustration of the PD1 LIRECAP assay. Equimolar concentrations of P and C aptamers were added to biospecimens, bound aptamers extracted and aptamers amplified together using a single set of primers and quantified using unique TaqMan colorimetric probes. The ratio of bound aptamers (C:P ratio) was determined. (B) Raji-PDL1 and Raji-Null cells were combined in varying ratios (0%–100% PDL1 positivity) and co-cultured with Jurkat-PD1 cells at a 1:1 Raji:Jurkat ratio. Jurkat-PD1 T cell inhibition mediated through the PD1 signaling was quantified as the percentage decrease in luciferase activity relative to the 0% Raji-PDL1 control in co-culture conditioned medium. After supernatant was collected, cell pellets were formalin-fixed and analyzed using the PD1 LIRECAP assay to assess the C:P ratio. (C) Raji-PDL1 cells correlate with inhibition of Jurkat-PD1 T cell activation (n=2). (D) Increasing the percentage of Raji-PDL1 increases the C:P ratio as determined by the PD1 LIRECAP assay (n=2). (E) C:P ratio as determined by the PD1 LIRECAP assay correlates with PD1 signaling-mediated T cell inhibition in the Raji-Jurkat co-culture system (n=2).

Jurkat-PD1 cells were cultured with various ratios of Raji-PDL1 and Raji-null cells (schema in Fig. 3B and Supplementary Table S3) to produce samples with varying PD1 saturation by PDL1. Increasing the fraction of Raji cells that expressed PDL1 in the coculture led to Jurkat-PD1 TCR inhibition and reduced luciferase activity (R^2^ = 0.962; p<0.0001) (Fig 3C). After signaling was determined, cocultured cells were pelleted, fixed in formalin and analyzed using the PD1 LIRECAP assay. Increasing the fraction of Raji cells expressing PDL1 in the co-culture increased in the C:P ratio (R^2^ = 0.832; p< 0.001) (Fig. 3D). There was a strong correlation between the C:P ratio and signaling inhibition (R^2^ = 0.810; p<0.0001) (Fig. 3E). These data indicate that the C:P ratio as determined by the PD1 LIRECAP assay correlates with PD1 signaling, and that the PD1 LIRECAP assay can be performed on formalin fixed cells.

### Evaluation of Formalin-fixed paraffin-embedded (FFPE) cell pellets

Formalin fixed cell pellets from mixtures of Jurkat and Raji cells were embedded in paraffin using standard clinical protocol. Tissue sections were cut from the resulting FFPE blocks and the PD1 LIRECAP assay and PLA performed (Fig. 4A). The C:P ratio determined by the PD1 LIRECAP assay correlated with increasing fraction of Raji-PDL1 cells in the FFPE samples (R^2^ = 0.975; p=0.013) (Fig. 4B). PD1-PDL1 proximity was measured using the Naveni PD1/PD-L1 Atto647N PLA kit (Navinci)[4]. PLA staining was consistent with the PD1 LIRECAP results (Fig. 4C) and revealed a higher percent of cells showing PLA positivity when greater numbers of Raji-PDL1 were present in the co-culture (R^2^ = 0.808; p<0.0001) (Fig. 4D). There was a strong correlation between PLA positivity and C:P ratio (R2=0.763; p<0.0001) (Fig. 4E).

**Fig. 4:**
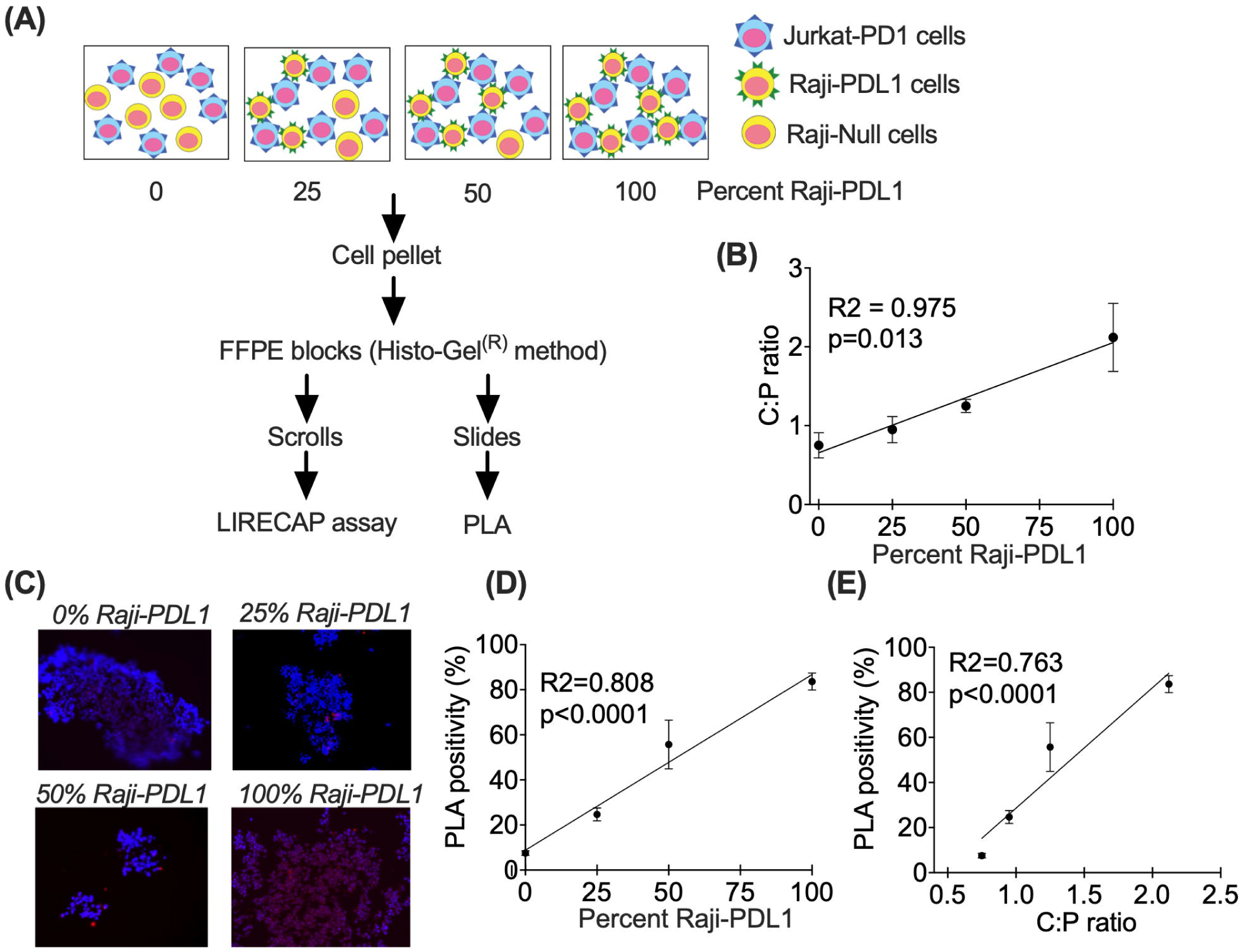
Correlation between PD1 LIRECAP assay (C:P aptamer ratio) and PD1-PDL1 proximity in FFPE biospecimens from Jurkat-Raji co-culture system. (A) Schematic illustration of the preparation and analysis of FFPE sections from the Jurkat-Raji co-culture system to assess the correlation between the PD1 LIRECAP assay and PD1-PDL1 proximity. (B) C:P ratio as measured by PD1 LIRECAP assay correlates with PD1 saturation by PDL1 (n=2) (C) Representative images from PD1-PDL1 PLA of FFPE from cell pellets (Atto647N fluorescence) (n=2). (D) Quantification of PLA positivity per cell, expressed as percent positive cells (n=2). (E) PD1-PDL1 PLA correlates with C:P aptamer ratio, as measured by PD1 LIRECAP assay (n=2).

### Technical reproducibility, intratumoral heterogeneity and interpatient variability in PD1 saturation by PDL1

Six human sarcoma FFPE biospecimens with T cell infiltration were selected for initial evaluation of clinical biospecimens (Supplementary Table S4). Three were identified as PDL1^low^ (Specimens A, B and C) and three as PDL1^high^ (Specimens D, E and F). To evaluate the technical reproducibility of the assay on tumor specimens, biospecimens were deparaffinized, macerated, divided into multiple aliquots and the PD1 LIRECAP assay run separately on matched aliquots. Linear mixed effects regression of the repeated C:P ratios used to estimate intraclass correlation coefficient (ICC) indicated a high level of technical reproducibility (ICC = 0.943; 95% CI = 0.814 – 0.978) (Fig. 5A), including when each aliquot from the same section was assayed at separate time points (ICC = 0.849; 95% CI = 0.085 – 0.972) (Fig. 5B). A similar level of technical reproducibility was seen when sequential FFPE sections from Raji-Jurkat cell pellets (as described in Fig.4) were evaluated (Overall ICC = 0.954; 95% CI = 0.764 – 0.987).

**Fig. 5:**
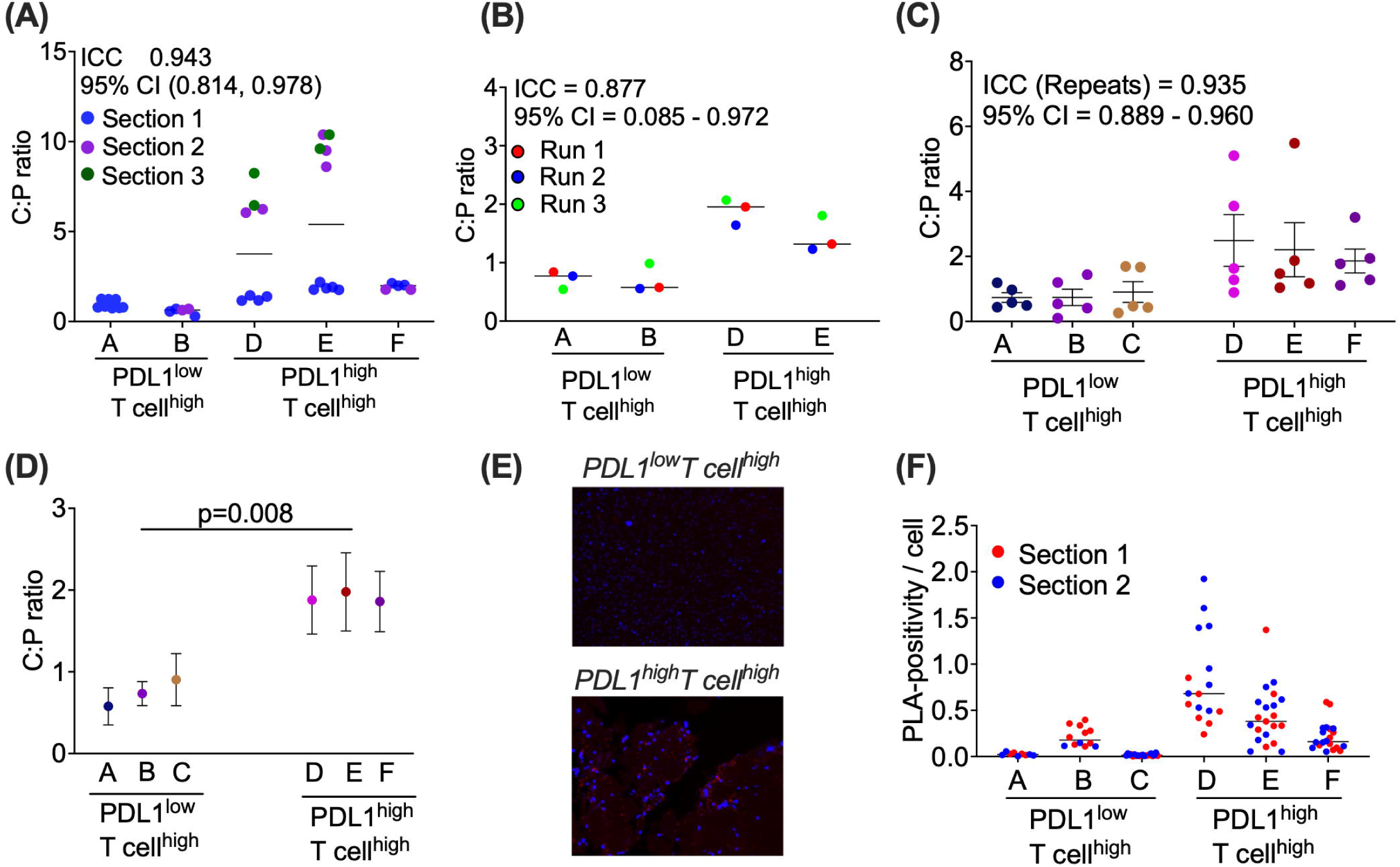
Quantification of PD1 saturation on human clinical sarcoma FFPE biospecimens. Sections were obtained from six archived sarcoma biospecimens and categorized as PDL1^high^T cell^high^ and PDL1^low^T cell^high^. Sections representing different regions within the tumors were deparaffinized and evaluated by the PD1 LIRECAP assay and PLA. (A) Technical reproducibility of repeated measurements done on multiple aliquots per section (repeated measurements from same scroll are marked with same color). (n = 5 tumors) (B) Technical reproducibility of the PD1 LIRECAP assay when assay was performed at different times/runs. (n = 4 tumors) (C) Intratumoral heterogeneity of C:P ratio (n = 5 sections per tumor representing different regions) (D) Interpatient variability of C:P ratio. (E) Representative images from PLA on same biospecimens. (E) Quantification of PLA positivity per cell in different regions of the tumor confirming intratumoral heterogeneity in PD1 and PDL1 proximity. (n = 2 sections per tumor).

To assess intratumoral heterogeneity, multiple tissue sections were analyzed from each tissue block. There was considerable heterogeneity in the C:P ratio when different tissue samples from the same block were evaluated. Linear mixed effect regression analysis indicated that 93.5% of the observed C:P ratio variability is caused by the biological intratumoral variability in the PD1 saturation by PDL1 (ICC = 0.935; 95% CI = 0.889 – 0.960) (Fig. 5C). When the average C:P ratio was determined for each tumor, there was a higher C:P ratio in the PDL1^high^ biospecimens compared to PDL1^low^ biospecimens (p=0.008; Fig. 5D). PLA staining was done in parallel, indicating brighter PLA staining in PDL1^high^ than PDL1^low^ biospecimens. In corroboration with PD1 LIRECAP assay (Fig. 5E), PLA positivity also demonstrated high intratumoral heterogeneity (Covariance: (PDL1^low^: A = 63.76%, B = 49.10%, and C = 68.29%; PDL1^low^: D = 59.03%, E = 70.33%, and F = 70.09) (Fig. 5F).

### Developing a standard curve to convert C:P ratio to PD1 saturation by PDL1

The specificity of the aptamers used in the PD1 LIRECAP assay is relative, not absolute, with some cross-reactivity. In the PD1 LIRECAP assay, this cross-reactivity is normalized by quantifying the ratio of bound C18 to bound P17 (C:P ratio) concurrently in a single qPCR assay. The C:P ratio data can then be converted to a metric of PD1 saturation by PDL1, termed as the PD1-PDL1 Saturation Index, using a standard curve. To develop such a standard curve, various concentrations of recombinant PDL1 were added to PD1-coated beads and formalin fixed as reported in Fig. 2 above. PD1 saturation was calculated using the quadratic formula (see methods)[26]. As illustrated in Fig. 6, the C:P ratio correlated well with the calculated PD1 saturation by PDL1 (R^2^ = 0.956; p = 0.0001). Such standard curve, when run in parallel with the test biospecimens, can be used to quantitatively analyze the PD1 saturation index of clinical samples.

**Fig. 6:**
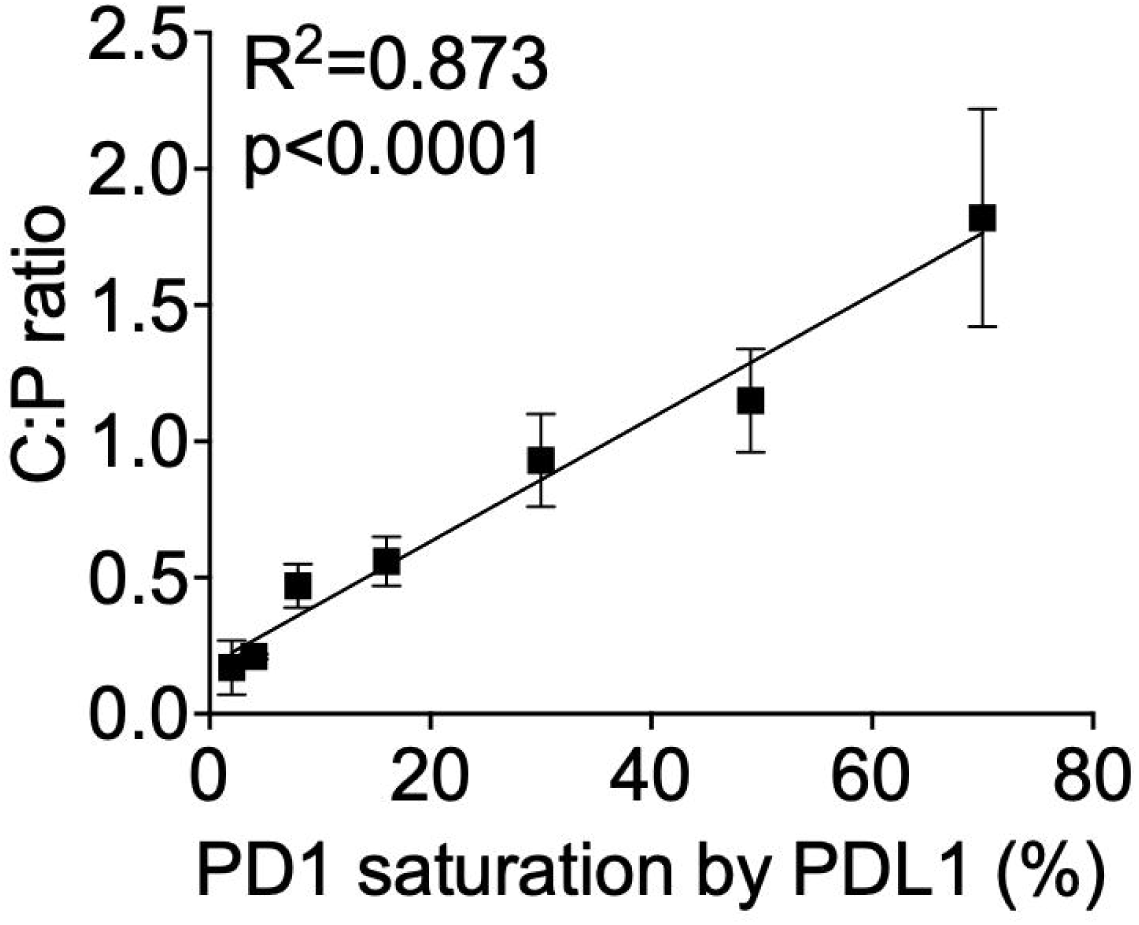
Standard curve to convert C:P ratio to PD1 Saturation Index. Beads with known PD1 saturation by PDL1 were produced using recombinant human PD1 and PDL1, formalin-fixed, washed and evaluated by the PD1 LIRECAP assay to determine C:P ratios. C:P ratios were plotted against the PD1 saturation by PDL1 to create a standard curve. Data shown are the mean and SEM of two independent experiments.

## Discussion

Biomarkers currently used to predict the likelihood of response to PD1 blockade therapy are suboptimal[28, 29]. They are of modest value for patients with some types of cancer and of no clear value for many others [15, 30], [31]. Better biomarkers would be of considerable patient benefit[8]. Quantifying saturation of PD1 by PDL1 in the tumor has the potential to be such a biomarker based on the assumption that anti-PD1-based therapy is more likely to be effective in those patients with high PD1 saturation by PDL1 in the tumor. This concept is supported by studies utilizing PD1/PDL1 proximity assays, such as FRET or PLA to predict response[32], [20] however such assays are challenging to implement on a large scale[4]. The LIRECAP assay platform was designed to overcome this limitation by leveraging unique properties of RNA aptamers. Importantly, this platform is particularly well suited to quantify the saturation of PD1 by PDL1 using equipment and techniques available in standard laboratories.

RNA aptamers can fold into stems and loops and bind to target antigens in a manner analogous to antibodies. The SELEX methodology allows for identification of aptamers that have different binding specificities to targets in their native confirmation, i.e., without the need for antigen processing or presentation. Therefore, RNA aptamers can be identified that recognize epitopes on individual molecules or interacting molecules (such as receptor-ligand pairs). Aptamers chosen from a single library can have different specificities but are of the same length and have the same 5’ and 3’ ends. Leveraging these properties, the LIRECAP assay utilizes one aptamer that preferentially binds to unoccupied PD1 and one aptamer that preferentially binds to the PD1-PDL1 complex. Such aptamers were added together to a biospecimen, bound aptamers extracted and amplified using standard two-color qPCR with a single set of primers. The specificity of the aptamers using in the LIRECAP assay for unoccupied PD1 or the PD1-PDL1 complex is relative, not absolute. Such cross-reactivity, which complicates other assay platforms, is overcome in the LIRECAP assay by expanding and quantifying both aptamers simultaneously in a single assay and assessing their ratio. This results in an extraordinary degree of internal control. The C:P aptamer ratio can then be converted to a PD1 Saturation Index through use of a standard curve.

Here we describe the development and preliminary validation of the PD1 LIRECAP assay. Candidate aptamer sequences that preferentially bound to unoccupied PD1 or to the PD1-PDL1 Complex were identified using a modification of the SELEX process. Screening also included evaluation of aptamer binding to formalin-fixed target proteins. This fixation stabilized the receptor-ligand complex, thereby assuring sample manipulation or the aptamers themselves doesn’t disrupt the receptor-ligand interaction and it confirmed the selected aptamers can detect targets in FFPE biospecimens without an additional antigen retrieval processing. Determination of PD1 saturation using the PD1 LIRECAP assay reflected PD1 signaling and PD1-PDL1 proximity, in a cell model.

The PD1 LIRECAP assay was evaluated in a pilot study with archived clinical FFPE sarcoma tissue that allowed for evaluation of technical reproducibility, intratumoral heterogeneity and interpatient variability. Technically, the PD1 LIRECAP assay was highly reproducible with consistent C:P ratios on individual biospecimens that were evaluated at different times. There was considerable intratumoral heterogeneity in PD1 saturation by PDL1 when different regions of a given biospecimen were assessed, a feature that was also seen when PLA was used to determine PD1-PDL1 proximity in the same tissue. This is consistent with the literature indicating spatial variability of PD1 and PDL1 within individual tumors across various cancer types[17, 18, 33–35]. Despite intratumoral heterogeneity in PD1 saturation, there were significant differences between tumors when the average PD1 saturations were compared among biospecimens. Biospecimens with high PDL1 expression had higher PD1 saturation than those with low PDL1 expression.

Several key questions related to the PD1 LIRECAP assay remain. Intratumoral heterogeneity represents a challenge for all tissue-based prognostic biomarkers. We do not yet know whether PD1 saturation correlates with clinical outcome in response to PD1 blockade. Such studies are ongoing. Not all PD1 and PDL1 is found on the cell surface with studies indicating that PDL1 can be found in soluble form, in exosomes and even in the nucleus[17]. The potential impact of non-cell surface PD1 or PDL1 on the PD1 LIRECAP assay is unclear. The ability of the assay to quantify PD1 saturation by PDL2 requires further delineation. Ongoing optimization studies should clarify how these factors impact on interpretation of the assay’s results and clinical applicability. Despite these remaining questions, the PD1 LIRECAP assay has significant promise as a clinical and a research tool. The assay can be performed using standard biospecimens, equipment and expertise readily available in most clinical and research laboratories. Thus, there should be no major hurdles to optimizing and validating the assay for clinical use should it prove to be clinically valuable and could provide valuable biological insight.

The studies reported here focused on a two-aptamer LIRECAP assay with one aptamer preferentially binding to PD1 and the other to the PD1-PDL1 complex. Modifications to this assay under development include a third aptamer preferentially binding to unoccupied PDL1 and even a fourth to a control house-keeping protein. Such an approach would allow for quantification of not only PD1 saturation by PDL1 on immune cells but also PDL1 saturation by PD1 on tumor tissue. We are continuing to make technical improvements to optimize the LIRECAP assay itself. Finally, ongoing studies are exploring how spatial analysis might be combined with the LIRECAP concept which could have significant clinical and research implications.

## Conclusions

Therapeutic agents targeting the PD1-PDL1 interaction are of great clinical value, however accurately predicting which patients are most likely to benefit is challenging. Improved predictive biomarkers for anti-PD1 therapy are clearly needed. Here, we describe development and preliminary evaluation of the PD1 LIRECAP assay, a novel assay capable of quantifying PD1 saturation by PDL1 in FFPE tumor tissue. Quantification of PD1 saturation by PDL1 as determined by the PD1 LIRECAP assay correlated closely with PD1-mediated signaling and PD1-PDL1 proximity. Analysis of clinical sarcoma FFPE biospecimens confirmed the assay is technically reproducible. There were significant differences in PD1 saturation by PDL1 between patients as well as considerable intratumoral heterogeneity. We conclude the assay is technically feasible, reproducible, and has the potential to serve as a superior predictive biomarker for PD1/PDL1-based therapy. Similar assays based on this platform could be used in other systems and settings to quantify interaction between two molecules.

## List of abbreviations

FFPE: Formalin-fixed, paraffin-embedded
LIRECAP: Ligand-Receptor Complex-binding Aptamers
PD1: Programmed cell death protein 1
PDL1: Programmed cell death protein ligand 1
PDL2: Programmed cell death protein ligand 2
PLA: Proximity Ligation assay
RNA: Ribonucleic acid
RT-qPCR: Reverse-transcriptase-quantitative polymerase chain reaction
SELEX: Systematic evolution of ligands by Exponential enrichment
TCR: T cell receptor

## Declarations

### Authors’ contributions

SV – Designed/planned/executed the study, interpreted the data and wrote the manuscript

CY – Executed Proximity ligation assay experiments and wrote the manuscript

NY – Preliminary characterization of Jurkat-Raji co-culture and wrote the manuscript

BJS – Analyzed the data and wrote of the manuscript

GJW – Provided overall direction, designed/planned the study, interpreted the data and wrote the manuscript

## Acknowledgements

The authors would like to acknowledge use of the Biospecimens and Molecular Epidemiology Core, the University of Iowa Central Microscopy Research Facility and the Genomics Division, core resources, University of Iowa Holden Comprehensive Cancer Center, University of Iowa Vice President for Research and the University of Iowa Carver College of Medicine.

