## Supplementary Information for "Quantifying PD1 saturation by PDL1 in tumor tissue using a novel RNA aptamer-based assay"

**Supplementary Figures:**

***Supplementary Fig. S1: Number of unique aptamer sequences during SELEX***

Data displaying a reduction in the number of unique RNA aptamer sequences with each round of SELEX, indicating progressive enrichment as rounds progressed.

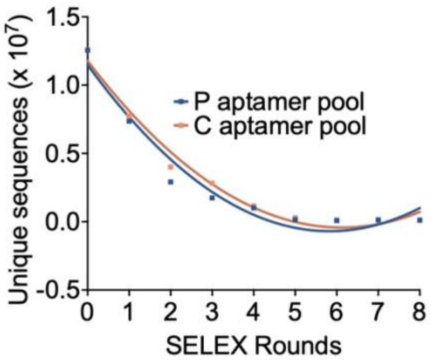


***Supplementary Fig. S2: Cross-reactivity of aptamers with PDL2 and PD1-PDL2 complex***

(A) P17 and (B) C18 aptamers display a preferential binding pattern towards PD1 and PD1-PDL2 complex, respectively, like the binding pattern with PD1-PDL1 complex, suggesting a structural similarity between the complexes suggesting the C aptamer binds to both the PD1-PDL1 Complex and the PD1-PDL2 Complex.

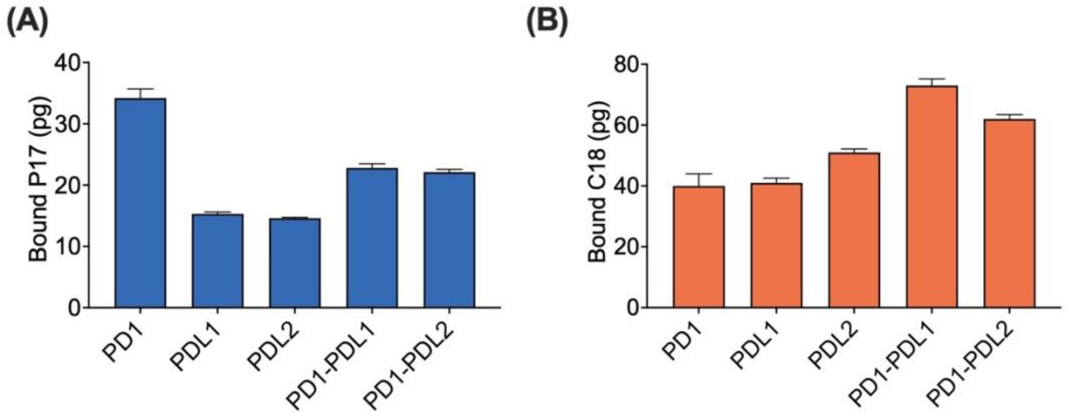

***Supplementary Fig. S3: Specificity of aptamers and probes for PD1 LIRECAP assay***

(A) P17 (blue) and C18 (orange) aptamers do not compete for binding to PD1 or PD1-PDL1 Complex and do not cross-block each other when added together in the PD1 LIRECAP assay. (B) TaqMan probe designed for P17 show high specificity to P17 aptamer, not C18 aptamer. (C) TaqMan probe designed for C18 show high specificity to C18 aptamer, not P17 aptamer.

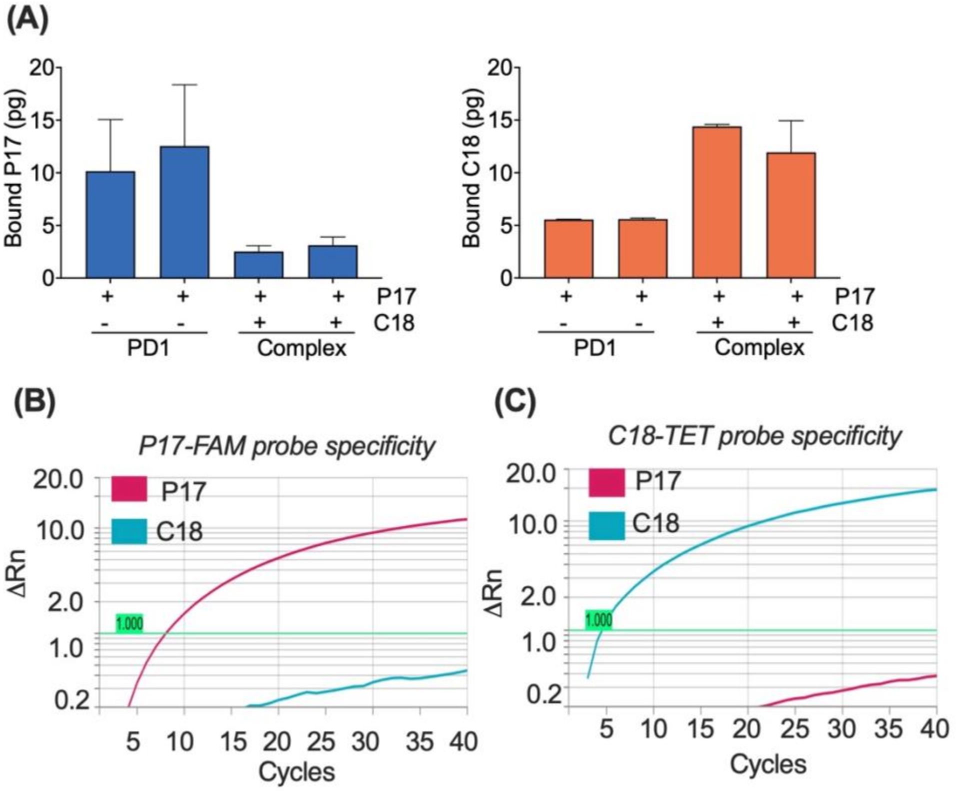

**Supplementary tables:**

***Supplementary Table S1: SELEX binding conditions***

<table>
<tr><th>SELEX rounds</th><th>Preclearing*</th><th>Post-clearing**</th><th>[RNA]</th><th>[Target protein]***</th><th>Binding time</th></tr>
<tr><td>1 & 2</td><td>200 nM His-HSA + human IgG1</td><td>N/A</td><td>2 μm</td><td>200 nM</td><td>120 min</td></tr>
<tr><td>3 & 4</td><td>200 nM His-HSA + human IgG1</td><td>N/A</td><td>2 μm</td><td>200 nM</td><td>60 min</td></tr>
<tr><td>5 & 6</td><td>200 nM His-HSA + human IgG1</td><td rowspan="4">1. 100 nM PDL1 for post-clearing PD1 SELEX<br>2. 100 nM PD1 followed by PDL1 for post-clearing Complex SELEX</td><td>1 μm</td><td>200 nM</td><td>45 min</td></tr>
<tr><td>7</td><td>300 nM His-HSA + His-UbQ + human IgG1</td><td>500 nM</td><td>50 nM</td><td>45 min</td></tr>
<tr><td>8</td><td>300 nM His-HSA + His-UbQ + human IgG1</td><td>500 nM</td><td>50 nM</td><td>30 min</td></tr>
<tr><td>9</td><td>500 nM His-HSA + His-UbQ + human IgG1</td><td>100 nM</td><td>25 nM</td><td>15 min</td></tr>
</table>

* Preclearing of RNA pool was performed before binding with target proteins to remove aptamers non-specifically binding to tags (6X His or Fc) and proteins

** Post-clearing of RNA pool was performed after the binding step with target proteins to remove aptamers that cross-reacted between PD1 and PDL1

*** Target proteins concentrations provided are based on the PD1 protein used for PD1 SELEX. For Complex SELEX, PDL1 was added to PD1-coated beads at enough concentration to create at least 75% PD1 saturation.[24]

***Supplementary Table S2: Fold enrichment of selected candidates from PD1 SELEX and Complex SELEX over successive SELEX rounds***

| *Aptamer name* | *Fold enrichment in respective SELEX (normalized to Rd 0)* | | | | |
|---|---|---|---|---|---|
| | *Rd 1* | *Rd 3* | *Rd 5* | *Rd 7* | *Rd 8* |
| P10 | 0.4 | 31.8 | 42.2 | 283.5 | 1413.3 |
| P17 | 0.3 | 6.8 | 22.6 | 152.7 | 661.5 |
| P150 | 1.0 | 174.5 | 73.3 | 218.7 | 194.3 |

| | | | | | |
|---|---|---|---|---|---|
| C18 | 0.4 | 1.3 | 134.9 | 541.8 | 1244.1 |
| C52 | 0.5 | 7.2 | 99.2 | 697.4 | 768.4 |
| C113 | 1.0 | 0 | 98.3 | 320.9 | 240.5 |

***Supplementary Table S3: Co-culturing Jurkat-PD1 and Raji-Null cells to obtain varying ranges of PD1 saturation by PDL1***

| *PDL1 to PD1 (%)* | *Jurkat-PD1* | *Raji-Null* | *Raji-PDL1* |
|---|---|---|---|
| *0* | *100* | *100* | *0* |
| *6.25* | *100* | *93.75* | *6.25* |
| *12.5* | *100* | *87.5* | *12.5* |
| *25* | *100* | *75* | *25* |
| *50* | *100* | *50* | *50* |
| *100* | *100* | *0* | *100* |

***Supplementary Table S4: Immunological characteristics of Human FFPE sarcoma biospecimens***

| | **Sample Name** | **IHC data** | | |
|---|---|---|---|---|
| | | PDL1 | Immune cells | CD3 |
| T cell$^{high}$PDL1$^{low}$ | A | - | + | + |
| | B | - | + | + |
| | C | - | + | + |
| T cell$^{high}$PDL1$^{high}$ | D | + | + | + |
| | E | + | + | + |
| | F | + | + | + |
